# Glycolytic Disruption Triggers Interorgan Signaling to Nonautonomously Restrict *Drosophila* Larval Growth

**DOI:** 10.1101/2024.06.06.597835

**Authors:** Madhulika Rai, Hongde Li, Robert A. Policastro, Gabriel E. Zentner, Travis Nemkov, Angelo D’Alessandro, Jason M. Tennessen

## Abstract

*Drosophila* larval growth requires efficient conversion of dietary nutrients into biomass. Lactate Dehydrogenase (Ldh) and Glycerol-3-phosphate dehydrogenase (Gpdh1) support larval biosynthetic metabolism by maintaining NAD^+^/NADH redox balance and promoting glycolytic flux. Consistent with the cooperative functions of Ldh and Gpdh1, the loss of both enzymes, but neither single enzyme, induces a developmental arrest. However, Ldh and Gpdh1 exhibit complex and often mutually exclusive expression patterns, suggesting that the *Gpdh1; Ldh* double mutant lethal phenotype could be mediated nonautonomously. Here we find that the developmental arrest displayed by the double mutants extends beyond simple metabolic disruption and instead stems, in part, from changes in systemic growth factor signaling. Specifically, we demonstrate that this synthetic lethality is linked to the upregulation of Upd3, a cytokine involved in the Jak/Stat signaling pathway. Moreover, we demonstrate that either loss of the Upd3 or dietary administration of the steroid hormone 20-hydroxyecdysone (20E) rescue the synthetic lethal phenotype of *Gpdh1; Ldh* double mutants. Together, these findings demonstrate that metabolic disruptions within a single tissue can nonautonomously modulate interorgan signaling to ensure synchronous developmental growth.

**SUMMARY STATEMENT:** We used the fruit fly *Drosophila melanogaster* to demonstrate that disruption of glycolysis within a single larval tissue alters systemic cytokine signaling and nonautonomously inhibits development of the entire animal.

## INTRODUCTION

Animal development requires the precise integration of environmental and metabolic cues with intrinsic growth signaling pathways (Miyazawa and Aulehla, 2018). Under ideal growth conditions, the metabolic pathways involved in central carbon metabolism efficiently convert dietary nutrients into the necessary biomass and energy to support rapid growth. Conversely, environmental stressors such as starvation, infection, and toxicant exposure necessitate metabolic reprogramming to maintain survival until conditions improve for growth (Bland, 2023, Deng and Kerppola, 2013, Koyama et al., 2020). The coordination between growth and metabolism under stressful environmental conditions, however, is complex as there is a need to monitor and coordinate metabolic flux across all tissues of the developing animal (Kim et al., 2021). When a single tissue encounters metabolic limitations, peripheral tissues must adjust their growth rates to prevent asynchronous development. This topic of interorgan metabolic communication has become a burgeoning area of research, with numerous studies examining how nutrient-sensing proteins and secreted signaling molecules coordinate growth across multiple tissues (Rajan and Perrimon, 2011, Droujinine and Perrimon, 2016, Castillo-Armengol et al., 2019). Nevertheless, the specific metabolic signals that trigger interorgan growth signaling are only beginning to emerge (Baker and Rutter, 2023, Wang and Lei, 2018, Figlia et al., 2020).

The fruit fly *Drosophila melanogaster* is a powerful system for exploring how metabolism is coordinated with developmental growth (Gillette et al., 2021). The ∼200-fold increase in body mass that occurs during the four days of *Drosophila* larval development represents an appealing model for exploring metabolic mechanisms that convert nutrients into biomass and energy. In this regard, previous studies revealed that this impressive growth rate is supported by a coordinated increase in the expression of genes involved in glycolysis, the pentose phosphate pathway, as well as *Ldh* (Rechsteiner, 1970, Tennessen et al., 2011, Tennessen et al., 2014). The resulting larval metabolic program exhibits hallmark features of aerobic glycolysis – a specialized form of carbohydrate metabolism that is ideally suited for rapid biomass production. Considering that several types of cancer cells also rely on aerobic glycolysis for growth and survival (Vander Heiden et al., 2009), *Drosophila* larvae present a compelling model to study this biosynthetic state.

We previously demonstrated that the larval metabolic program requires the enzymes Ldh and Gpdh1, which cooperatively regulate NAD^+^/NADH redox balance and promote glycolytic flux (Li et al., 2019). Intriguingly, while the loss of either single enzyme has minimal effect on overall larval growth, *Gpdh1; Ldh* double mutants experience a developmental arrest, suggesting a functional redundancy, where each enzyme partially compensates for the loss of the other (Li et al., 2019). However, neither Ldh nor Gpdh1 are ubiquitously expressed in larvae (Rechsteiner, 1970, Li et al., 2019, Rai et al., 2024), raising questions as to how these enzymes serve compensatory roles if not expressed in a strictly overlapping pattern. Here we address this question by examining the tissue-specific functions of both enzymes.

Our analysis of Gpdh1 and Ldh expression during larval development confirms previous observations that these enzymes are expressed in a complex and often mutually exclusive expression pattern (Rechsteiner, 1970). Moreover, we demonstrate that the loss of both enzymes within a single tissue inhibits larval growth in a nonautonomous manner. Subsequent RNA-seq analysis of *Gpdh1; Ldh* double mutants reveals altered expression of several secreted signaling molecules, including increased expression of the cytokine Upd3, which was previously shown to regulate systemic larval growth in response to cellular stress (Romao et al., 2021). Using a genetic approach, we find that *upd3; Gpdh1; Ldh* triple mutants can survive larval development and eventually eclose into adults, indicating that increased Upd3 expression serves a significant role in the synthetic lethal phenotype of *Gpdh1; Ldh* double mutants. Finally, we demonstrate that the steroid hormone 20E also plays a role in this phenotype, as a dietary supplement of 20E also allows *Gpdh1; Ldh* double mutants to complete larval development. Overall, our findings reveal that the developmental delay and lethal phenotype associated with *Gpdh1; Ldh* double mutants are not simply the result of metabolic failure but rather stem, in part, from changes in systemic growth signaling.

## RESULTS AND DISCUSSIONS

### Ldh and Gpdh1 influence larval tissue growth in a nonautonomous manner

To better understand how Ldh and Gpdh1 cooperatively promote *Drosophila* larval growth, we examined the spatial expression pattern of both enzymes. Our analysis revealed that Ldh and Gpdh1 are often expressed in a mutually exclusive manner. In the Malpighian tubules, for example, *Ldh* is expressed in the stellate cells while *Gpdh1* is primarily expressed in principal cells (Figure 1A-A’’’). Similarly, cells of the larval midgut tend to express high levels of either Ldh or Gpdh1, but rarely do we observe a cell expressing both enzymes (Figure 1B-B’’’, S1A-D). This trend is also apparent in the central nervous system, where although both *Ldh* and *Gpdh1* are co-expressed in many cells the CNS, *Gpdh1*, but not *Ldh*, is expressed in neural stem cells and the prothoracic gland (Figure 1C-D’’’ and S1E-H). Finally, both the fat body and salivary glands express Gpdh1 but not Ldh (Figure 1A-A’’’, S1I-P), a result that is consistent with previous observations. In fact, the only tissue where we consistently observed uniform and overlapping expression of both enzymes was in the larval body wall muscle (Figure 1E-E’’’).

**Figure 1.**
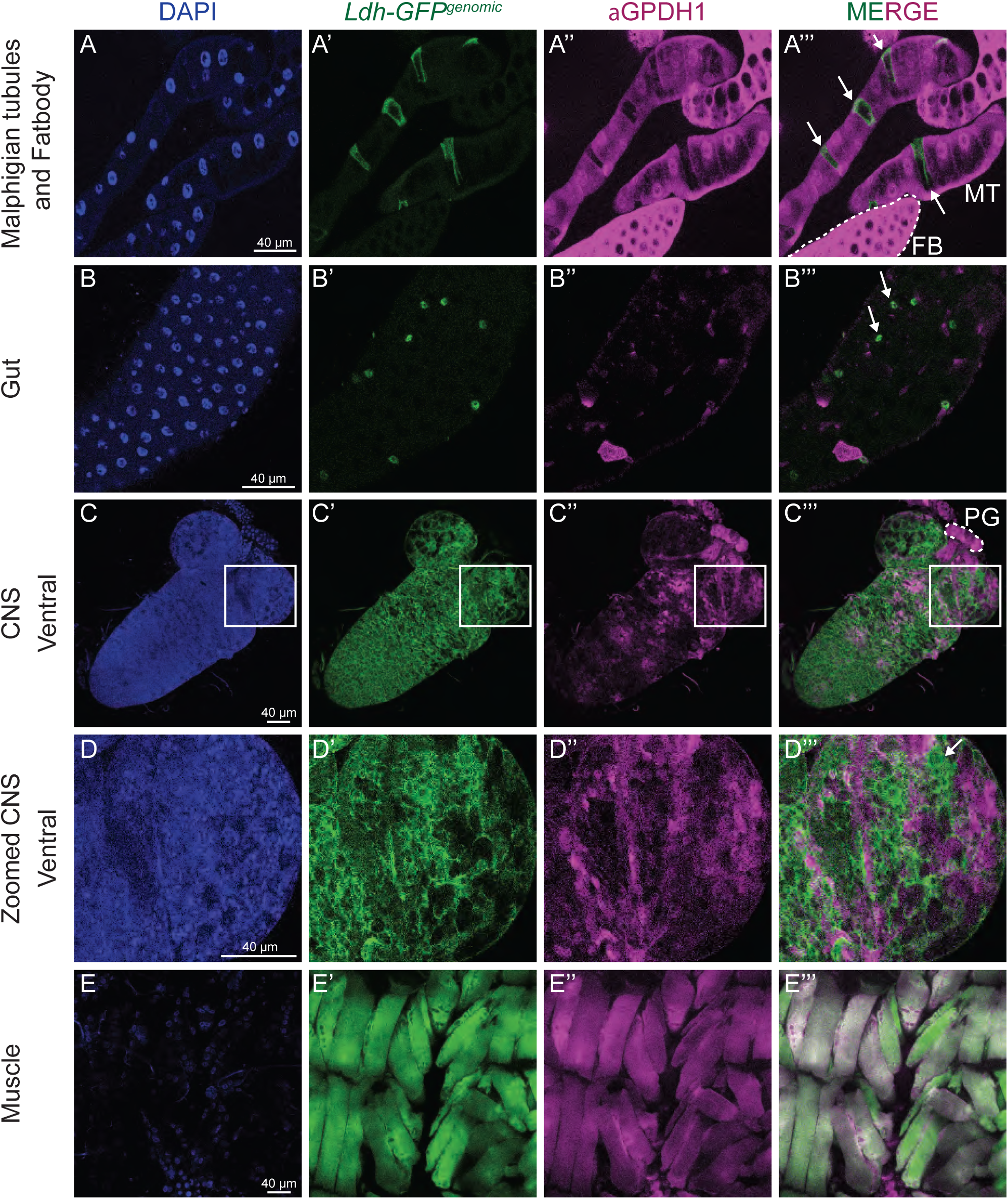
*Ldh* and *Gpdh1* expression patterns are complex and non-strictly overlapping. Representative confocal images of second instar larval tissues expressing *Ldh-GFP^Genomic^* and immuno-stained with αGpdh1 antibody. DAPI is shown in blue, Ldh-GFP and Gpdh1 are represented in green and magenta, respectively. The rightmost panel displays the merged images of Ldh-GFP and Gpdh1 staining. (A-A’’’) Malpighian tubules, (B-B’’’) gut, (C-C’’’) ventral side of CNS, (D-D’’’) magnification of the outlined region of interest in (C-C’’’), and (E-E’’’) muscles. Arrows denote non-overlapping Ldh expression. The scale bar represents 40 μM. The scale bar in (A) applies to all panels.

The complex *Ldh* and *Gpdh1* expression patterns raise the question as to how these enzymes cooperatively regulate larval metabolism if they are not present in the same cells and tissues. One explanation is that loss of one enzyme alters the spatial expression pattern of the other; however, we find no evidence to support this hypothesis. For example, Ldh expression is nearly undetectable in wild-type salivary glands and is not increased in *Gpdh1* mutant salivary glands (Figure S2A-G). Similarly, salivary gland Gpdh1 expression is not further elevated in *Ldh* mutants (Figure S2H-N). We also observe no increase in Ldh expression within the fat body of *Gpdh1* mutants (Figure S3A-G), although we do see a slight increase in Gpdh1 expression within *Ldh* mutant fat bodies (Figure S3 H-N). These results are also consistent with a previous observation that Gpdh1 levels are not increased in *Ldh* mutant clones within the larval CNS (Li et al., 2019).

Our findings suggest that the *Gpdh1; Ldh* double mutant growth defects are not simply the combined result of metabolic dysfunction within individual cells, but rather stem from changes in interorgan signaling. We tested this hypothesis by using RNAi to deplete *Ldh* expression in the muscle (*Mef2R-GAL4)* and fat body (*r4-GAL4*) of *Gpdh1* mutant larvae. With both drivers, loss of the two enzymes within a single tissue significantly reduced larval growth (Figure 2A, S4A) and slowed development as determined by time to pupation (Figure 2B, S4B). Moreover, these developmental delays were apparent in tissues not expressing the *Ldh-RNAi* transgene – for example, the salivary glands of age-matched *Gpdh1; Mef2R-Ldh-RNAi* larvae were significantly smaller than those of either the wild-type control, *Gpdh1* mutant, or *Mef2R-Ldh-RNAi* animals (Figure 2C-F). Together, these results suggest that loss of both enzymes within a single tissue can globally influence larval growth and development.

**Figure 2.**
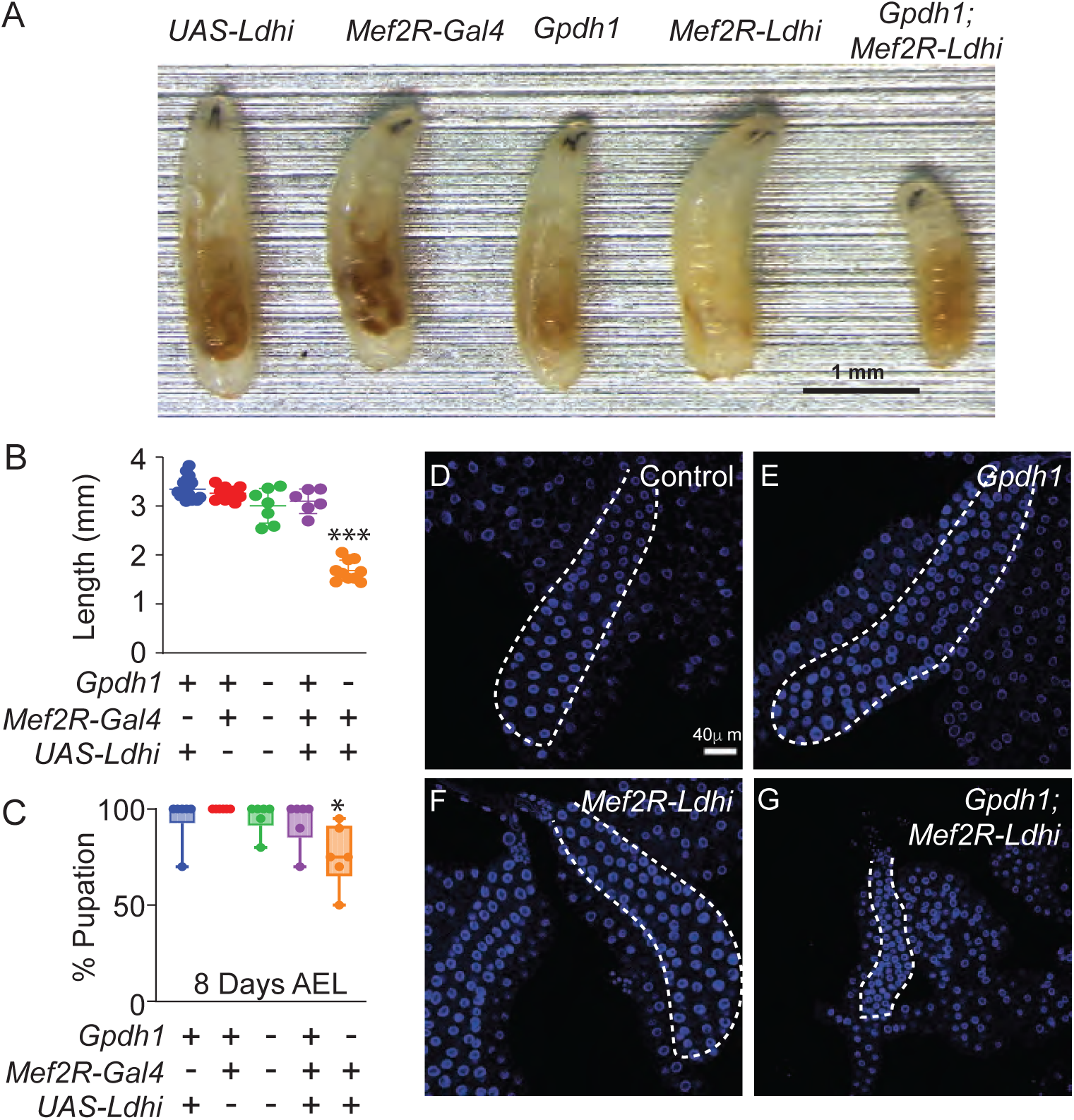
Tissue-specific loss of Ldh and Gpdh1 induces systemic growth defects. Growth and development of both control (*Mef2R-Gal4* and *UAS-Ldh-RNAi* strains) and mutant strains (*Gpdh1^A10/B18^* mutants, *Mef2R-Ldh-RNAi*, and *Gpdh1^A10/B18^; Mef2R-Ldh-RNAi*) were monitored throughout larval development. (A) Representative images of L2 larvae (60 hr AEL) from the indicated genotypes. The scale bar represents 1 mm. (B-C) Quantification of (B) larval length and (C) time to pupation from the indicated genotypes. All experiments were repeated a minimum of three times. n≥5 biological replicates for (B, C). Error bars represent standard deviation; \**P*<0.05, \*\*\**P*<0.0001. *P*-values were calculated using the Mann-Whitney test. (D-G) Representative images of salivary glands dissected from the indicated genotypes. DAPI is shown in blue. The scale bar represents 40 μM. The scale bar in (D) applies to (E).

### Upd3 is required to arrest larval growth in *Gpdh1; Ldh* double mutants

To better understand how Ldh and Gpdh1 nonautonomously regulate larval development, we used RNA-seq to analyze gene expression in *Ldh^16/17^*mutants, *Gpdh1^A10/B18^* mutants, and *Gpdh1^A10/B18^; Ldh^16/17^* double mutants relative to genetically-matched heterozygous controls (Table S1). While our approach identified several genes that were significantly changed in both single mutants (Figure 3A-B and S5), for the purpose of this analysis we focused on those genes that were only altered in double mutants (Table S2). Among the genes that were significantly down- or up-regulated in *Gpdh1^A10/B18^; Ldh^16/17^* double mutants but not the single mutants, we identified four secreted factors that were previously shown to systemically regulate larval growth and metabolism: *peptidoglycan recognition protein SC2* (*PGRP-SC2*; FBgn0043575), *peptidoglycan recognition protein LA*, (*PGRP-LA*; FBgn0035975), *dawdle* (*daw*; FBgn0031461), and *unpaired 3* (*upd3*; FBgn0053542) (Droujinine and Perrimon, 2016, Musselman et al., 2018, Romao et al., 2021). While each of these molecules is known to function in interorgan communication and could contribute towards regulating the growth of *Gpdh1; Ldh* double mutants, we decided to focus on the cytokine Upd3 for the remainder of this study.

**Figure 3.**
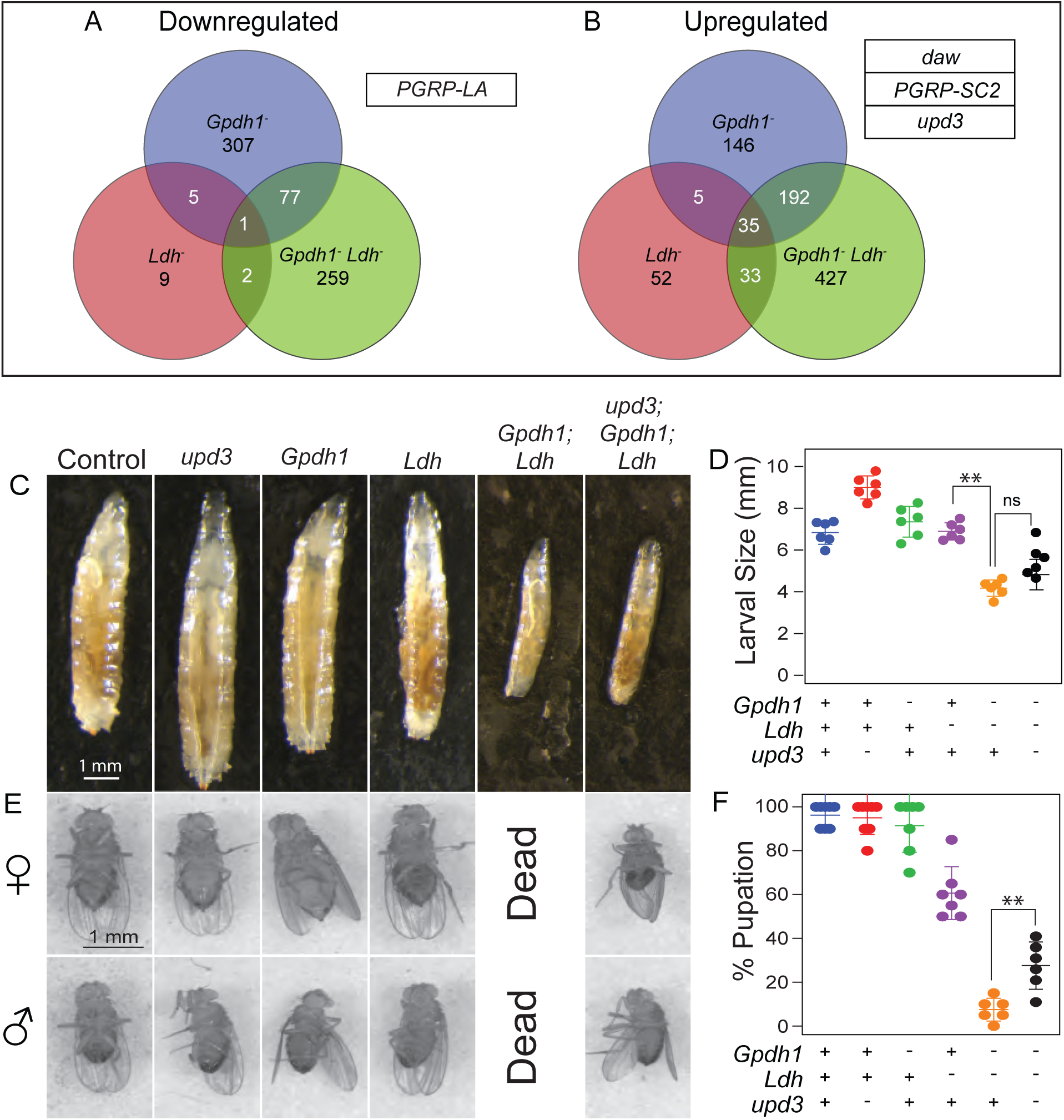
An *upd3* mutation suppresses the synthetic lethal phenotype of *Gpdh1; Ldh* double mutants. (A-B) Venn diagrams showing the overlap between the number of genes that were either (A) downregulated or (B) upregulated in either single mutant (*Gpdh1^A10/B18^* and *Ldh^16/17^*) as well as the double mutant (*Gpdh1^A10/B18^; Ldh^16/17^*) relative to the respective heterozygous controls strains. Associated tables in (A, B) list secreted factors that are only dysregulated in the *Gpdh1^A10/B18^; Ldh^16/17^* double mutant. (C) Representative larval images of the indicated genotypes. The scale bars represent 1 mm and apply to all panels. (D) Quantification of the larval size for the indicated genotypes. (E) A significant number of male and female triple mutant pupae (*upd3^Δ^; Ldh^16/17^; Gpdh1^A10/B18^*) successfully eclosed whereas all *Gpdh1^A10/B18^; Ldh^16/17^*died during metamorphosis. The scale bar represents 1mm and applies to all panels. (F) The rate of eclosion was quantified for pupae of the indicated genotypes. Note that no eclosion was observed among vials containing *Gpdh1^A10/B18^; Ldh^16/17^* pupae. All experiments are repeated a minimum of three times. n≥5 biological replicates. Error bars represent standard deviation; \*\**P*<0.05. *P*-values were calculated by using a Mann-Whitney test.

Our RNA-seq analysis indicates that *upd3* expression is significantly elevated in *Gpdh1; Ldh* double mutants. Considering that Upd3 inhibits larval development in response to both environmental and intracellular stress (Droujinine and Perrimon, 2016, Romao et al., 2021, Yang et al., 2015), increased expression of this cytokine could serve a role in the *Gpdh1; Ldh* developmental arrest phenotype. We tested this possibility by generating *upd3^τι^; Gpdh1^A10/B18^; Ldh^16/17^* triple mutants, with the hypothesis that loss of Upd3 signaling in the double mutant background would suppress the larval arrest phenotype. Indeed, triple mutants completed larval development and entered metamorphosis at a significantly higher rate than the double mutants (Figure 3C-D). Moreover, while we never observed a *Gpdh1^A10/B18^; Ldh^16/17^*double mutant complete metamorphosis, up to 30% of triple mutant larvae survived to adulthood (Figure 3E-F). We would note, however, that *upd3^τ<^; Gpdh1^A10/B18^; Ldh^16/17^* triple mutants were significantly smaller than controls, sick, and died within 2-3 days of eclosion, indicating that not all aspects of the *Gpdh1^A10/B18^; Ldh^16/17^* double mutant phenotype are regulated by Upd3.

To better understand how the *upd3* mutations suppress the *Gpdh1^A10/B18^; Ldh^16/17^* larval arrest phenotype, we examined both double and triple mutant larvae using a semi-targeted metabolomics approach (Table S3). Our analysis revealed no significant metabolic differences between the two strains (Figure S6). Notably, the levels of both lactate and glycerol-3-phosphate were unchanged in the triple mutant as compared with double mutant larvae (Figure S6B-C), indicating that the *upd3* mutation does not restore glycolytic metabolism in *Gpdh1^A10/B18^; Ldh^16/17^* double mutants. Overall, these results suggest that *Gpdh1^A10/B18^; Ldh^16/17^* double mutants do not simply die from metabolic dysfunction but rather experience a larval arrest due, in part, to increased Upd3 signaling.

### Ecdysone feeding is sufficient to suppress the *Gpdh1; Ldh* larval lethal phenotype

To further explore the connection between Upd3 and the *Gpdh1^A10/B18^; Ldh^16/17^* larval arrest phenotype, we used a *10XStat92E-GFP* reporter to identify those tissues with increased JAK/STAT signaling (Yang et al., 2015, Ekas et al., 2006). Consistent with Upd3 serving a key role in limiting systemic growth, *Gpdh1^A10/B18^; Ldh^16/17^* double mutants exhibited widespread *10XStat92E-GFP* reporter expression when compared with either the heterozygous control strain, *Ldh^16/17^* single mutant, or *Gpdh1^A10/B18^*single mutant. In this regard, we observed increased *10XStat92E-GFP* expression in the fat body (Figure S7A-D), salivary glands (Figure S7E-H), muscles (Figure S7I-L), and CNS (Figure 4A-D). Intriguingly, we also observed significantly increased *10XStat92E-GFP* expression in the prothoracic gland (PG) of double mutants relative to the single mutants (Figure 4A-D), although, we would note that *Gpdh1^A10/B18^* single mutants displayed elevated *10XStat92E-GFP* expression relative to either the *Ldh^16/17^* single mutant or heterozygous control strain (Figure 4B).

**Figure 4.**
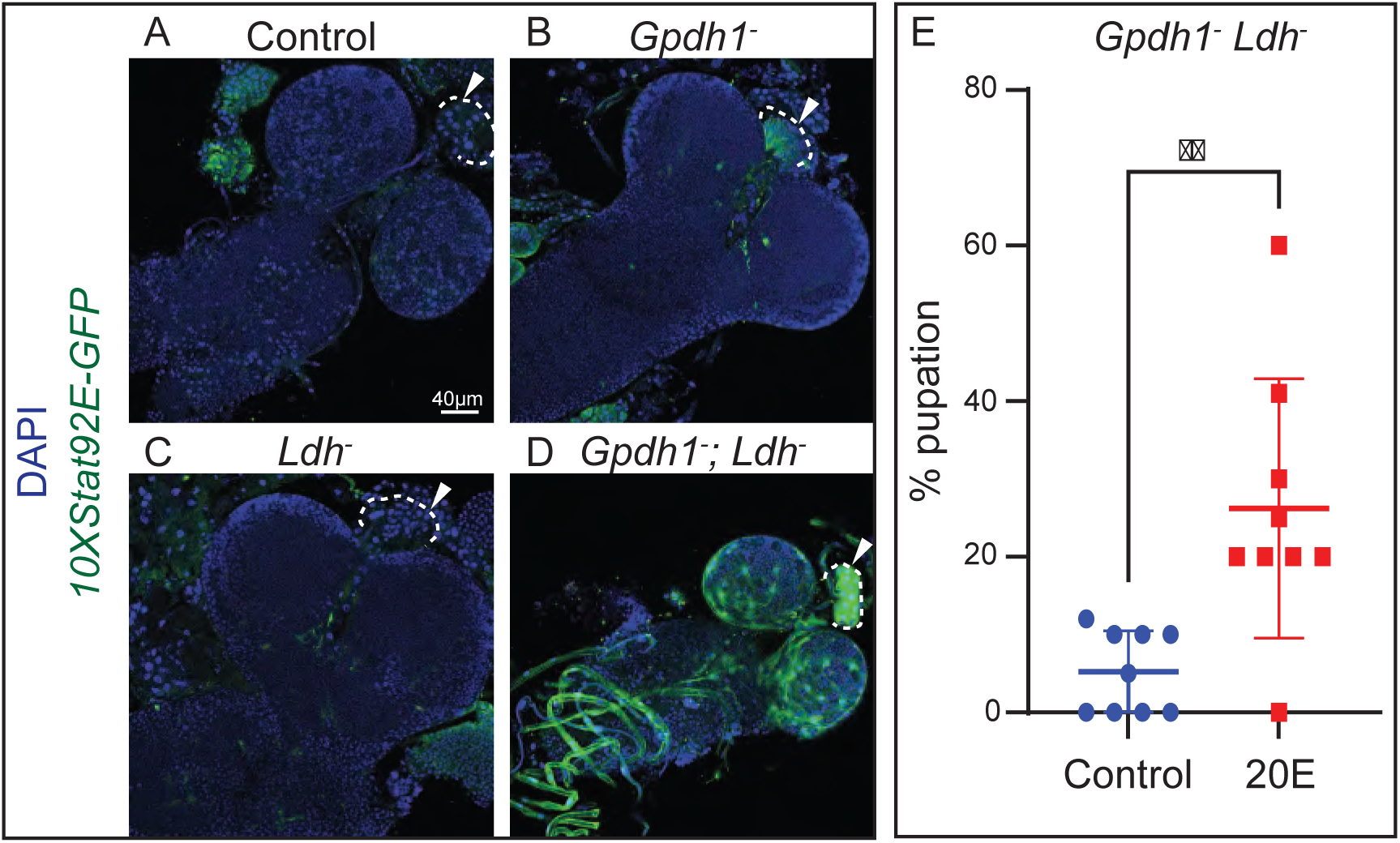
Dietary supplementation with 20E suppresses the synthetic lethal phenotype of *Gpdh1; Ldh* double mutants. (A-D) Representative confocal images of *Stat-GFP* expression in the CNS of *Gpdh1^A10/B18^; Ldh^16/17^* double mutants as compared to the heterozygous control (*Gpdh1^A10/+^; Ldh^16/+^*) and single mutant strains (*Gpdh1^A10/B18^*and *Ldh^16/17^*). DAPI is shown in blue and *Stat-GFP* expression in green. The scale bar in (A) represents 40 μM and applies to (B-D). (E) A graph illustrating the percent of *Gpdh1^A10/B18^; Ldh^16/17^* double mutant that pupariate when raised on yeast-molasses agar that contains either ecdysone or the solvent (ethanol) control. All experiments are repeated a minimum of three times. n=9 biological replicates. Error bars represent standard deviation. \*\**P*<0.05. *P* value was calculated using a Mann-Whitney test.

A previous study of tumorous wing discs demonstrated that Upd3 signaling can suppress ecdysone production in the PG and inhibit larval development (Romao et al., 2021). Thus, our observation that *10XStat92E-GFP* expression is elevated within the *Gpdh1^A10/B18^; Ldh^16/17^* PG raises the possibility that the double mutant larval arrest phenotype stems, in part, from decreased ecdysone signaling. Consistent with this model, we found that the addition of 20-hydroxyecdysone (20E; the active form of ecdysone) to the larval food significantly increased the pupariation rate of *Gpdh1^A10/B18^; Ldh^16/17^* double mutants (Figure 4E). Overall, these results indicate that changes in 20E signaling contribute to the *Gpdh1^A10/B18^; Ldh^16/17^* double mutant synthetic lethal phenotype.

### Redefining the developmental functions of “Housekeeping Genes”

Our studies unexpectedly reveal that *Drosophila* larval development can tolerate loss of two key enzymes that regulate cytosolic NAD^+^/NADH balance. These findings, while surprising, are consistent with classic studies of *COX5A* (also known as *tenured*), which revealed that loss of this electron transport chain subunit induces a specific eye development phenotype (Mandal et al., 2005). The defects induced by *COX5A* mutations are not simply the result of defects in ATP synthesis, but rather stem from the activation of a pathway involving AMPK and p53, which eliminates Cyclin E and induces a cell cycle arrest (Mandal et al., 2005). These foundational observations were subsequently validated by both follow-up studies and a large-scale genetic screen, which demonstrated that depletion of many genes that encode electron transport chain subunits induce distinct defects in eye development (Liao et al., 2006, Pletcher et al., 2019). Our findings are built upon these earlier studies and again demonstrate that *Drosophila* larvae experiencing severe metabolic disruptions don’t simply die due to metabolic stress, but instead arrest development through the activity of specific growth signaling pathways. Specifically, our findings suggest that loss of both Gpdh1 and Ldh activate a previously described feedforward loop, in which elevated Upd3 signaling repressed 20E signaling (Romao et al., 2021).

Conceptually, our findings offer an interesting mechanistic rationale explaining how animals in the wild might cope with developmental delays as a function of exposure to severe metabolic stressors. While factors such as starvation and hypoxia inevitably induce cell autonomous responses involving the entire organism (Schito and Rey, 2018), animals also encounter environmental insults that affect individual tissues, including bacterial and parasitic infections, toxins that target specific organs, and nutrient imbalances that impinge upon cell-specific metabolic bottlenecks (Koyama et al., 2020). Animal development has evolved to identify these tissue-specific metabolic stresses and produce signals from effected cells that are subsequently amplified and recognized by unaffected tissues, thus allowing development to be delayed in a coordinate manner. For example, *Drosophila* hemocytes respond to parasitic wasp infections by inducing a cell intrinsic increase in glycolytic metabolism that not only facilitates an effective immune response but also produces an adenosine signal that remodels peripheral metabolism (Bajgar et al. 2015, Bajgar and Dolezal, 2018).

Our findings that Upd3 and 20E are part of a coordinated response to tissue-specific disruption of glycolytic flux provide new insights into the coordination between metabolism and developmental growth. Upd3 has emerged as a common stress-induced cytokine that communicates cell-specific information with peripheral tissues (Woodcock et al., 2015, Gera et al., 2022), and our finding reinforces a previously proposed model in which elevated Upd3 signaling induces a feedforward loop that inhibits 20E signaling and delays *Drosophila* development (Romao et al., 2021). In this regard, the involvement of a steroid hormone is an important feature of the regulatory mechanism, as it allows for a single tissue to indirectly influence the developmental progression of entire animal. Moving forward, the key question emerges of how metabolic stress regulates Upd3 signaling? Both individual metabolites as well end products of cell-specific metabolic dysfunction (e.g., redox imbalance and ROS accumulation) represent potential signals that could trigger Upd3 signaling as well as related modes of interorgan communication. Regardless of the answer, our study highlights that core metabolic enzymes are not simple products of “housekeeping genes,” but rather serve dynamic cellular functions that are fully integrated into the signaling pathways that regulate multicellular growth and development.

## METHODS

### *Drosophila melanogaster* husbandry and genetic analysis

Fly stocks were maintained at 25°C on Bloomington Drosophila Stock Center (BDSC) food. Larvae were raised and collected as previously described (Li and Tennessen, 2017). Unless noted, *Ldh* mutant larvae harbored a trans-heterozygous combination of *Ldh^16^* (RRID:BDSC_94698) and *Ldh^17^* (RRID:BDSC_94699) (Li and Tennessen, 2017) and *Gpdh1* mutant larvae harbored a trans-heterozygous combination of *Gpdh1^A10^* (RRID:BDSC_94702) and *Gpdh1^B18^* (RRID:BDSC_94703) (Li et al., 2019). The *upd3* mutant strain was obtained from the BDSC (RRID:BDSC_55728). The *Ldh-GFP^Genomic^*stock (RRID:BDSC_94704) used for colocalization analysis of Ldh and Gpdh1 was previously described (Bawa et al., 2020; Rai et al., 2024). The *w[1118]; P{w[+mC]=10XStat92E-GFP}2* stock was obtained from the BDSC (RRID:BDSC_26198) and recombined with *Ldh^16^* to visualize STAT expression levels in the tissues of (*Gpdh1^A10/B18^*) double mutant larvae. Tissue specific knockdown of Ldh was conducted using transgenes that express GAL4 in either muscle (P{w[+mC]=GAL4-Mef2.R}3; RRID:BDSC_27390) or fat body (P{w[+mC]=r4-GAL4}3; RRID:BDSC_33832). RNAi experiments were conducted using transgenes that targeted *Ldh* expression (RRID:BDSC_33640). Each of these Gal4 lines were placed into a *Gpdh1^B18^*mutant background and crossed to *w^1118^*; *Gpdh1^A10^*; *UAS-Ldh-RNAi* to attain the tissue specific knockdown. Flybase was used as an essential reference tool throughout this study (Gramates et al., 2022, Ozturk-Colak et al., 2024).

### Immunofluorescence

Larval tissues were dissected in 1X phosphate buffer saline (PBS; pH7.0) and fixed with 4% paraformaldehyde in PBS for 30 minutes at room temperature. Fixed samples were subsequently washed once with PBS and twice with 0.3% PBT (1x PBS with Triton X-100) for 10 mins per wash.

For GFP antibody staining, fixed tissues were incubated with goat serum blocking buffer (4% Goat Serum, 0.3% PBT) for one hour at RT and stained overnight at 4 °C with primary antibody Rabbit anti-GFP diluted 1:500 (#A11122 Thermo Fisher). Samples were washed three times using 0.3% PBT and stained with secondary antibody Alexa Fluor 488 Goat anti-Rabbit diluted 1:1000 (#R37116; Thermo Fisher) for either 4 hrs at room temperature or overnight at 4 °C. Stained tissues were washed with 0.3% PBT, immersed in DAPI (0.5µg/ml 1X PBS) for 15 mins and then mounted with a vector shield with DAPI (Vector Laboratories; H-1200-10).

Ldh protein expression was visualized using an anti-Ldh antibody (Boster bio DZ41222) as previously described (Rai et al., 2024). For larval tissues staining with anti-Gpdh1 antibody (Boster Bio DZ41223), the same protocol was used as for anti-Ldh staining, except that the anti-Gpdh1 antibody was diluted 1:100.

### Mounting and imaging

All the stained tissues were mounted using VECTASHIELD containing DAPI (Vector Laboratories; H-1200-10). Multiple Z-scans of individual tissues were acquired using the Leica SP8 confocal microscope in the Light Microscopy Imaging Center at Indiana University, Bloomington. Unless noted, figure panels contain a representative Z-scan. For presenting the Z stack in a 2D image, the Z projection tool of Fiji was used, with maximum projection.

Quantification of Gpdh1 and Ldh antibody staining was conducted by measuring the mean intensities for Gpdh1, Ldh and DAPI in the region of interest (ROI) per image, using Fiji. Gpdh1 and Ldh mean intensities were normalized to the mean DAPI intensity for the same ROI per image and plotted by using GraphPad Prism v10.1. Statistical analysis was conducted using the Mann-Whitney test.

### Imaging and quantification of larval size

Third instar larvae were collected and imaged using Leica MZ 10F microscope. The size of the larvae was measured by drawing a line from posterior to anterior end and noting down the total length by using Fiji. Data was plotted by using GraphPad Prism v10.1. Statistical analysis was conducted using the Mann-Whitney test.

### Adult imaging

Third instar wandering larvae were kept on a molasses agar plate with yeast paste. Starting on D9 after egg laying, pupae were checked for eclosion, and the newly eclosed adults were collected into a BDSC food vial. Adults were imaged by using Leica DFC 450.

### Percentage pupation

The pupation rate was calculated by placing 20 synchronized L1 larvae of each genotype on a molasses agar plate with yeast paste. Starting on day 4 after egg laying, plates were monitored every 24 hours for pupae.

### Sample collection and extraction for Metabolomics Analysis

Mutant and control larvae were collected at the mid-L2 stage (∼60 hours after egg-laying at 25°C) and extracted as previously described (Li and Tennessen, 2018). Briefly, collected larvae were placed into a prechilled 1.5 mL centrifuge tube on ice, washed with ice-cold water, and immediately frozen in liquid nitrogen. For extraction, larvae were transferred to pretared 1.4 mm bead tubes, the mass was measured using a Mettler Toledo XS105 balance, and the sample was returned to a liquid nitrogen bath prior to being stored at -80°C. Samples were subsequently homogenized in 0.8 mL of prechilled (-20 °C) 90% methanol containing 2 μg/mL succinic-d4 acid, for 30 seconds at 6.45 m/s using a bead mill homogenizer located in a 4°C temperature control room. The homogenized samples were incubated at -20 °C for 1 hour and then centrifuged at 20,000 × g for 5 min at 4°C. The resulting supernatant was sent for metabolomics analysis at the University of Colorado Anschutz Medical Campus, as previously described (Nemkov et al., 2019).

### Statistical analysis of metabolite data

The metabolomics dataset was analyzed using Metaboanalyst version 6.0 (Pang et al., 2024) with data normalized to sample mass and preprocessed using log normalization and Pareto scaling. Graphs for the relative levels of individual metabolites were plotted by using GraphPad Prism v10.1. Statistical analysis for these graphs were conducted using a Mann-Whitney test.

### Transcriptomic analysis

Quality control of reads performed with FastQC v0.11.9 (https://www.bioinformatics.babraham.ac.uk/projects/fastqc/). Read quantification to transcripts was accomplished with Salmon v1.4.0 (Patro et al., 2017) using the ENSEMBL D. melanogaster BDGP6.28 cDNA sequences file, and the softmasked toplevel ENSEMBL BDGP6.28 genome assembly as the decoy sequences. Reads were imported into R and aggregated into gene counts using tximport v1.18.0 (Soneson et al., 2015), and differential expression was performed using DESeq2 v1.30.0 (Love et al., 2014)

### Ecdysone feeding

Larvae were fed ecdysone in a modified molasses agar food. 115 mL of molasses and 24 grams of agar were added to 750 mL of boiling water on a stirring hot plate. The temperature was then reduced to 60°C and 5 grams of yeast was added to the media, at which point 2 mL of media was pipetted into a 35 mm petri dish. The low agar concentration resulted in a semi-solid media gained a semi-solid texture that was suitable for diffusion of a 20E stock solution.

A 30 mM stock solution of 20E (Sigma H5142) was prepared in 90% ethanol and 3.33 μL of stock solution was added to the modified molasses food, resulting in a final concentration of 50 μM (Ono, 2014; Koyama et al., 2014). Larvae of the tested genotypes were transferred to the 20E-supplemented food plates immediately after hatching and subsequently transferred to fresh 20E-supplemented plates every 24 hrs.

## ACKNOWLEDGEMENTS

We thank the Bloomington *Drosophila* Stock Center (NIH P40OD018537) for providing fly stocks, the *Drosophila* Genomics Resource Center (NIH 2P40OD010949) for genomic reagents, Flybase (NIH 5U41HG000739), and the Indiana University Light Microscopy Imaging Center. Thanks to Shefali Shefali for critical comments on the manuscript.

## COMPETING INTERESTS

No competing interests declared

## FUNDING

J.M.T. is supported by NIH 1R35GM119557.

## DATA AVAILABILITY

The RNA-seq data discussed in this publication have been deposited in NCBI’s Gene Expression Omnibus (Edgar *et al*., 2002) and are accessible through GEO Series accession number GSE119334 (https://www.ncbi.nlm.nih.gov/geo/query/acc.cgi?acc=GSE119334) and GSE121876 (https://www.ncbi.nlm.nih.gov/geo/query/acc.cgi?acc=GSE121876). Supplemental Tables S1-S3 are available at https://doi.org/10.6084/m9.figshare.25984402.v1.

## Supplementary

**Supplementary Figure 1. Ldh and Gpdh1 expression patterns in the CNS, fat body, salivary gland, and intestine.** Representative confocal images of second instar larval tissues expressing *Ldh-GFP^Genomic^* and immuno-stained with αGpdh1 antibody. DAPI is shown in blue, Ldh-GFP and Gpdh1 are represented in green and magenta, respectively. The rightmost panel displays the merged images of Ldh-GFP and Gpdh1 staining. (A-D) Dorsal side of CNS, (E-H) at body (I-L) salivary gland (M-P) gut. The scale bar in all the images represents 40 μM. The scale bar in (A) applies to (B-L) and the scale bar in (M) applies to (N,O,P).

**Supplementary Figure 2. Characterization of either Ldh or Gpdh1 expression in larval tissues following loss of the other enzyme.** Quantification of Ldh and Gpdh1 expression in the salivary glands of the heterozygous control strain (*Gpdh1^A10/+^; Ldh^16/+^*) and each single mutant strain (*Gpdh1^A10/B18^* and *Ldh^16/17^*). (A-F) Representative confocal images of (A-C) Ldh expression and (D-F) DAPI staining in all three genotypes. (G) Ldh staining was quantified in the salivary glands of all three genotypes. (H-M) Representative confocal images of (H-J) Gpdh1 expression and (K-M) DAPI staining in all three genotypes. (N) Gpdh1 staining was quantified in the salivary glands of all three genotypes. The scale bar in all images represents 40 μM. The scale bar in (A) applies to (B-F) and the scale bar in (H) applies to (I-M). (G, N) All experiments are repeated a minimum of three times. n=4 biological replicates. Error bars represent standard deviation. \*\**P*< 0.01. \**P*< 0.05. *P*-values were calculated by using the Mann-Whitney test.

**Supplementary Figure 3. Characterization of either Ldh or Gpdh1 expression in larval fat body following loss of the other enzyme.** Quantification of Ldh and Gpdh1 expression in the larval fat body of the heterozygous control strain (*Gpdh1^A10/+^; Ldh^16/+^*) and each single mutant strain (*Gpdh1^A10/B18^* and *Ldh^16/17^*). (A-F) Representative confocal images of (A-C) Ldh expression and (D-F) DAPI staining in all three genotypes. (G) Ldh staining was quantified in the fat body of all three genotypes. (H-M) Representative confocal images of (H-J) Gpdh1 expression and (K-M) DAPI staining in all three genotypes. (N) Gpdh1 staining was quantified in the larval fat body of all three genotypes. The scale bar in all images represents 40 μM. The scale bar in (A) applies to (B-F) and the scale bar in (H) applies to (I-M). (G, N) All experiments are repeated a minimum of three times. N=4 biological replicates. Error bars represent standard deviation. \*\**P* < 0.01. \**P* < 0.05. *P*-values were calculated by using the Mann-Whitney test.

**Supplementary Figure 4. Fat body expression of a *Ldh-RNAi* transgene in the fat body of *Gpdh1* mutants induces systemic growth defects.** Growth and development of both control (*r4-Gal4* and *UAS-Ldh-RNAi* strains) and mutant strains (*Gpdh1^A10/B18^*mutants, *r4-Ldh-RNAi*, and *Gpdh1^A10/B18^; r4-Ldh-RNAi*) were monitored throughout larval development. (A) Representative images of L2 larvae (60 hr AEL) from the indicated genotypes. The scale bar represents 1 mm. (B-C) Quantification of (B) larval length and (C) time to pupation of the indicated genotypes. (B, C) All experiments are repeated a minimum of three times. n≥6 biological replicates. Error bars represent standard deviation. \**P*<0.05 and \*\**P*<0.01 when compared with all other genotypes. *P*-values were calculated using the Mann-Whitney test.

**Supplementary Figure 5. RNA-seq analysis of *Ldh* mutants, *Gpdh1* mutants, and *Gpdh1, Ldh* double mutants.** Volcano plot depicting the transcriptomic profiles of (A) *Ldh* mutants (*Ldh^16/17^*), (B) *Gpdh1* mutants (*Gpdh1^A10/B18^*), and (C) double mutants (*Gpdh1^A10/B18^; Ldh^16/17^*) relative to the respective heterozygous control strains. n=3 biological replicates analyzed per genotype. Each sample contained 20 mid-L2 larvae. Vertical axis indicates -log10(FDR) and horizontal axis represents log(FC). The significantly upregulated genes are shown in yellow and downregulated are shown in black. FDR-fold discovery rate and FC-fold change.

**Supplementary Figure 6. *upd3; Gpdh1; Ldh* triple mutants exhibit a metabolomic profile that is indistinguishable from *Gpdh1; Ldh* double mutants.** (A) A volcano plot showing no significant difference between the metabolomes of *Gpdh1; Ldh* double mutants and *upd3; Gpdh1; Ldh* triple mutants. (B-C) Relative abundance of (B) lactate and (C) glycerol-3-phosphate in the indicated genotypes. n=6 biological replicates were collected from independent populations with 25 mid L2 larvae per sample. Experiment was repeated twice. Error bars represent standard deviation. No significant differences (ns) were observed in in double mutants as compared with the triple mutants. *P*-value was calculated by using the Mann-Whitney test. (D) Heatmap showing the top 25 metabolites rank order by significance among four genotypes - heterozygous control (HC), *upd3^Δ^* (U), *Gpdh1^A10/B18^; Ldh^16/17^* double mutant (DM) and the *upd3Δ; Gpdh1^A10/B18^; Ldh^16/17^* triple mutants (TM).

**Supplementary Figure 7. *Stat-GFP* expression is increased in larval tissues of *Gpdh1; Ldh* double mutants.** (A-L) Representative confocal images of (A-D) fat body, (E-H) salivary glands and (I-L) muscles showing *Stat-GFP* expression in control, *Gpdh1^A10/B18^, Ldh^16/17^* and *Gpdh1^A10/B18^; Ldh^16/17^* double mutants. The scale bar represents 40 μM. The scale bar in (A) applies to (B-L).

**Table S1.** RNA-seq analysis of *Gpdh1^A10/B18^* single mutant larvae relative to *Gpdh1^A10/+^* heterozygous controls, *Ldh^16/17^*single mutant larvae relative to the *Ldh^16/+^* heterozygous controls, and *Gpdh1^A10/B18^; Ldh^16/17^* double mutant larvae relative to *Gpdh1^A10/+^; Ldh^16/+^* heterozygous controls.

**Table S2.** A list of the genes that are significantly downregulated or upregulated in *Gpdh1^A10/B18^; Ldh^16/17^*double mutant larvae relative to *Gpdh1^A10/+^; Ldh^16/+^* heterozygous controls, but are expressed at normal levels in single mutant larvae.

**Table S3.** Metabolomic analysis of *Gpdh1^A10/+^; Ldh^16/+^* heterozygous controls (HC), *upd3^Δ^*single mutants (U), *Gpdh1^A10/B18^; Ldh^16/17^* double mutant (DM), and *upd3^Δ^; Gpdh1^A10/B18^; Ldh^16/17^*triple mutants (TM).

## Notes

https://www.ncbi.nlm.nih.gov/geo/query/acc.cgi?acc=GSE119334

https://www.ncbi.nlm.nih.gov/geo/query/acc.cgi?acc=GSE121876

https://doi.org/10.6084/m9.figshare.25984402.v1

### Competing Interest Statement

The authors have declared no competing interest.

### Summary of Updates

Added missing author - Hongde Li

