## Supplementary material for "Glycolytic Disruption Triggers Interorgan Signaling to Nonautonomously Restrict *Drosophila* Larval Growth": Figure S1

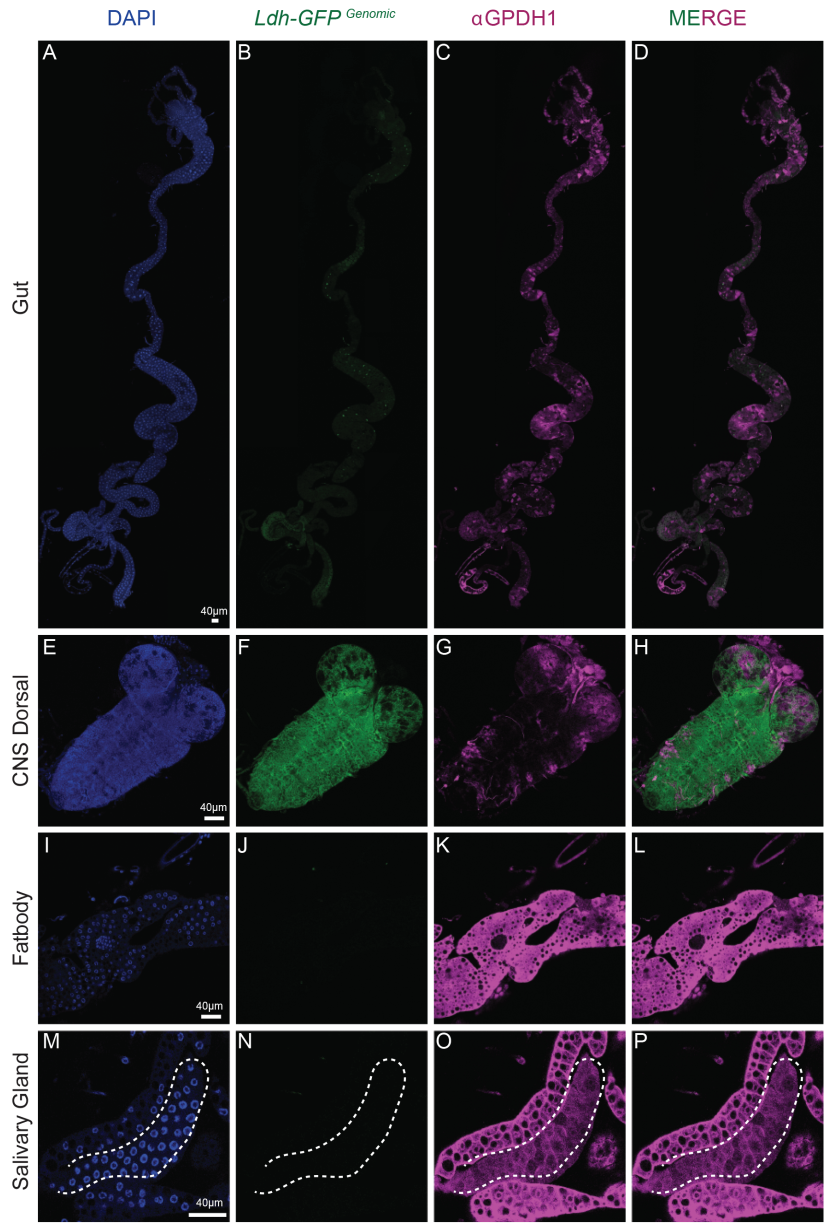


Figure S1

**Supplementary Figure 1. Ldh and Gpdh1 expression patterns in the CNS, fat body, salivary gland, and intestine.** Representative confocal images of second instar larval tissues expressing *Ldh-GFP*$^{Genomic}$ and immuno-stained with αGpdh1 antibody. DAPI is shown in blue, Ldh-GFP and Gpdh1 are represented in green and magenta, respectively. The rightmost panel displays the merged images of Ldh-GFP and Gpdh1 staining. (A-D) Dorsal side of CNS, (E-H) at body (I-L) salivary gland (M-P) gut. The scale bar in all the images represents 40 µM. The scale bar in (A) applies to (B-L) and the scale bar in (M) applies to (N,O,P).
