## Supplementary material for "Glycolytic Disruption Triggers Interorgan Signaling to Nonautonomously Restrict *Drosophila* Larval Growth": Figure S2

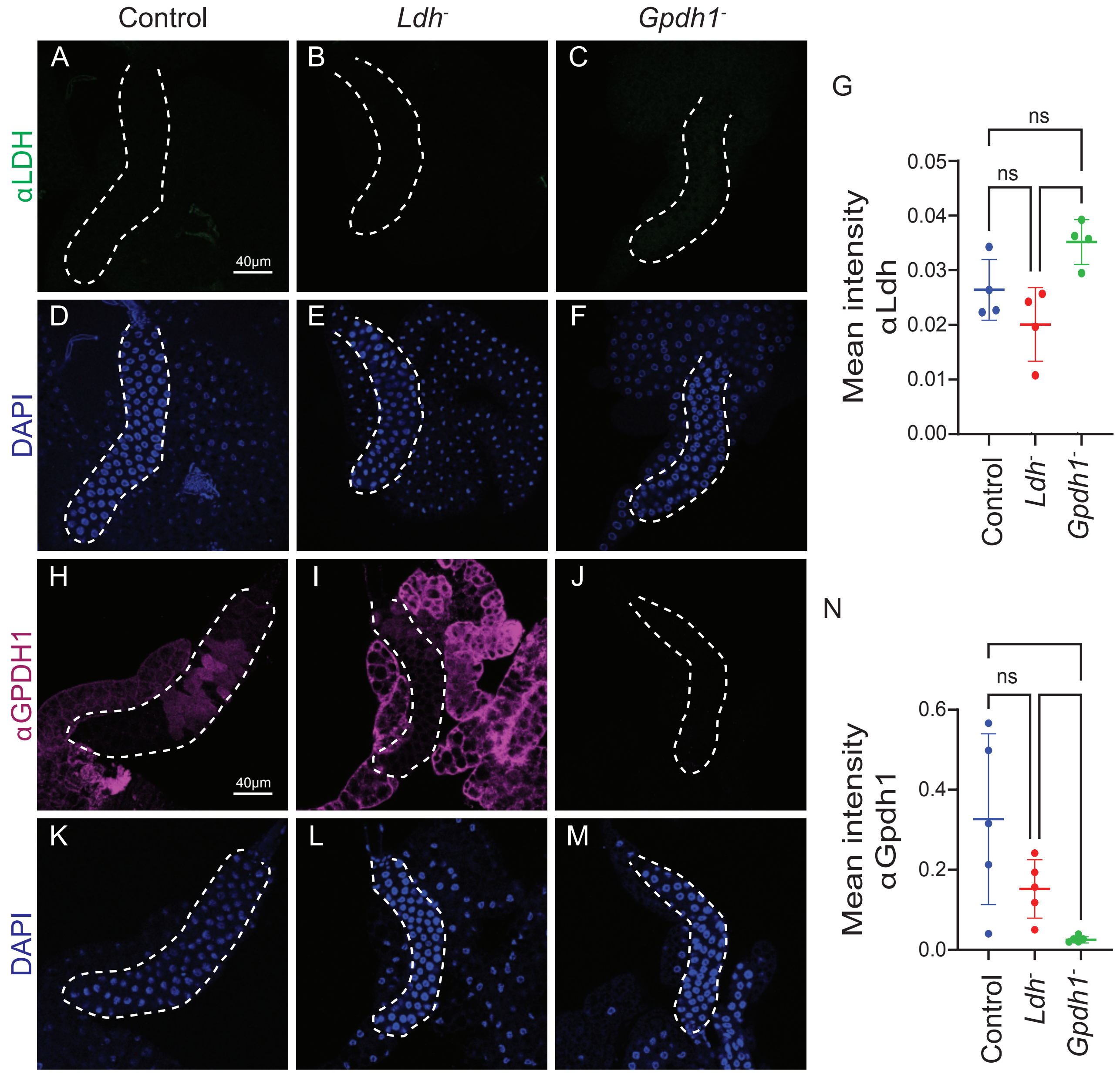


**Supplementary Figure 2. Characterization of either Ldh or Gpdh1 expression in larval tissues following loss of the other enzyme.** Quantification of Ldh and Gpdh1 expression in the salivary glands of the heterozygous control strain (*Gpdh1*$^{A10/+}$; *Ldh*$^{16/+}$) and each single mutant strain (*Gpdh1*$^{A10/B18}$ and *Ldh*$^{16/17}$). (A-F) Representative confocal images of (A-C) Ldh expression and (D-F) DAPI staining in all three genotypes. (G) Ldh staining was quantified in the salivary glands of all three genotypes. (H-M) Representative confocal images of (H-J) Gpdh1 expression and (K-M) DAPI staining in all three genotypes. (N) Gpdh1 staining was quantified in the salivary glands of all three genotypes. The scale bar in all images represents 40 μM. The scale bar in (A) applies to (B-F) and the scale bar in (H) applies to (I-M). (G, N) All experiments are repeated a minimum of three times. n=4 biological replicates. Error bars represent standard deviation. **$P$< 0.01. *$P$< 0.05. $P$-values were calculated by using the Mann-Whitney test.
