## Supplementary material for "Glycolytic Disruption Triggers Interorgan Signaling to Nonautonomously Restrict *Drosophila* Larval Growth": Figure S4

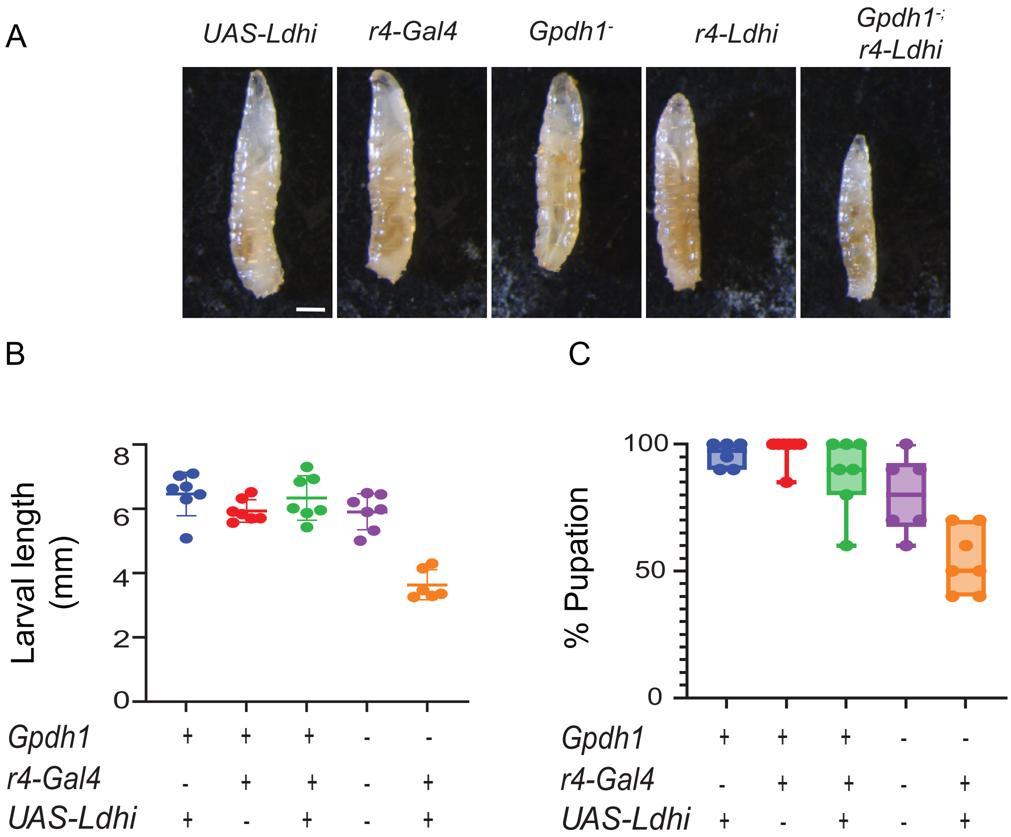


**Supplementary Figure 4. Fat body expression of a *Ldh-RNAi* transgene in the fat body of *Gpdh1* mutants induces systemic growth defects.** Growth and development of both control (*r4-Gal4* and *UAS-Ldh-RNAi* strains) and mutant strains (*Gpdh1*$^{A10/B18}$ mutants, *r4-Ldh-RNAi*, and *Gpdh1*$^{A10/B18}$; *r4-Ldh-RNAi*) were monitored throughout larval development. (A) Representative images of L2 larvae (60 hr AEL) from the indicated genotypes. The scale bar represents 1 mm. (B-C) Quantification of (B) larval length and (C) time to pupation of the indicated genotypes. (B, C) All experiments are repeated a minimum of three times. n≥6 biological replicates. Error bars represent standard deviation. *$P$<0.05 and **$P$<0.01 when compared with all other genotypes. $P$-values were calculated using the Mann-Whitney test.
