## Supplementary material for "Glycolytic Disruption Triggers Interorgan Signaling to Nonautonomously Restrict *Drosophila* Larval Growth": Figure S5

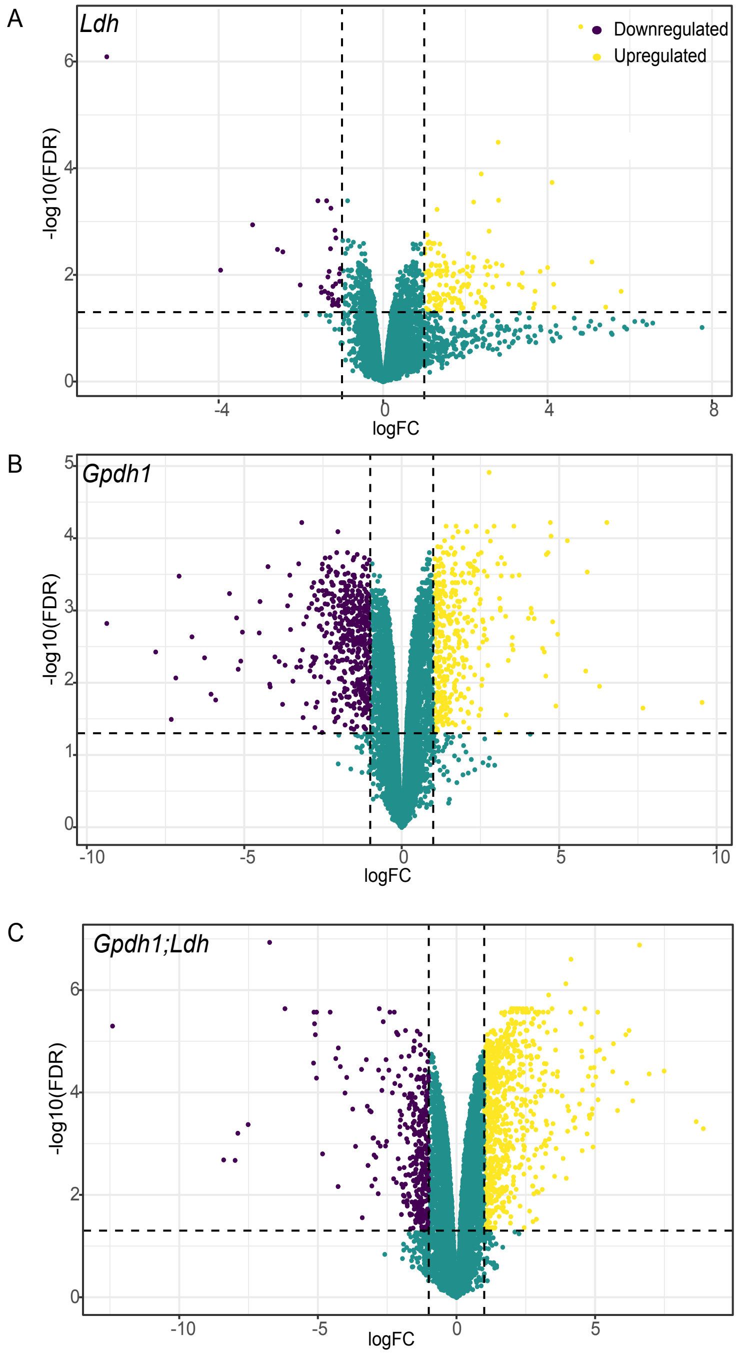


**Supplementary Figure 5. RNA-seq analysis of *Ldh* mutants, *Gpdh1* mutants, and *Gpdh1, Ldh* double mutants.** Volcano plot depicting the transcriptomic profiles of (A) *Ldh* mutants (*Ldh*$^{16/17}$), (B) *Gpdh1* mutants (*Gpdh1*$^{A10/B18}$), and (C) double mutants (*Gpdh1*$^{A10/B18}$; *Ldh*$^{16/17}$) relative to the respective heterozygous control strains. n=3 biological replicates analyzed per genotype. Each sample contained 20 mid-L2 larvae. Vertical axis indicates -log10(FDR) and horizontal axis represents log(FC). The significantly upregulated genes are shown in yellow and downregulated are shown in black. FDR- fold discovery rate and FC-fold change.
