## Supplementary material for "Glycolytic Disruption Triggers Interorgan Signaling to Nonautonomously Restrict *Drosophila* Larval Growth": Figure S6

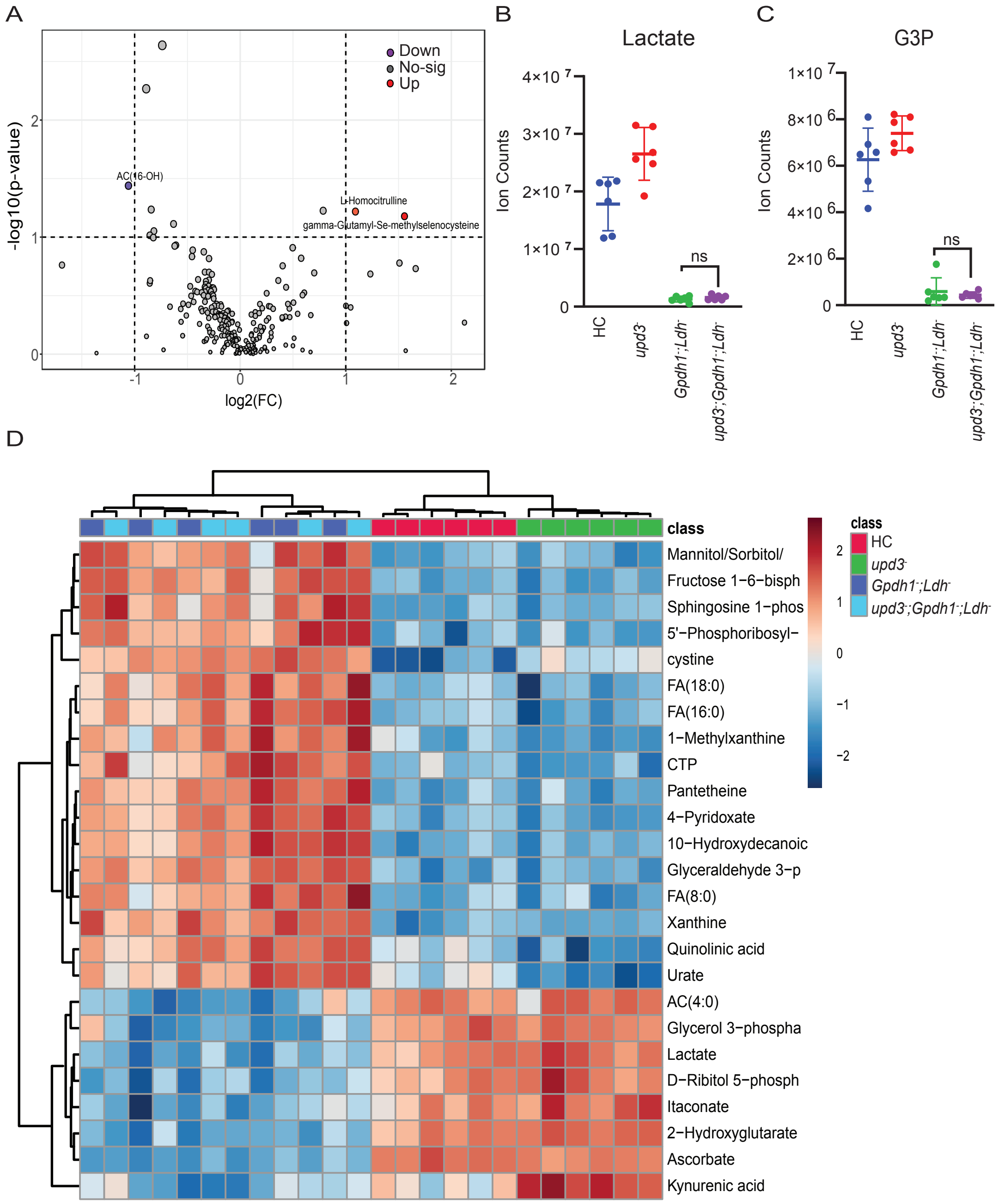


**Supplementary Figure 6. *upd3; Gpdh1; Ldh* triple mutants exhibit a metabolomic profile that is indistinguishable from *Gpdh1; Ldh* double mutants.** (A) A volcano plot showing no significant difference between the metabolomes of *Gpdh1; Ldh* double mutants and *upd3; Gpdh1; Ldh* triple mutants. (B-C) Relative abundance of (B) lactate and (C) glycerol-3-phosphate in the indicated genotypes. n=6 biological replicates were collected from independent populations with 25 mid L2 larvae per sample. Experiment was repeated twice. Error bars represent standard deviation. No significant differences (ns) were observed in in double mutants as compared with the triple mutants. *P*-value was calculated by using the Mann-Whitney test. (D) Heatmap showing the top 25 metabolites rank order by significance among four genotypes - heterozygous control (HC), *upd3*$^{\Delta}$ (U), *Gpdh1*$^{A10/B18}$; *Ldh*$^{16/17}$ double mutant (DM) and the *upd3Δ; Gpdh1*$^{A10/B18}$; *Ldh*$^{16/17}$ triple mutants (TM).
