## Supplementary material for "Glycolytic Disruption Triggers Interorgan Signaling to Nonautonomously Restrict *Drosophila* Larval Growth": Figure S7

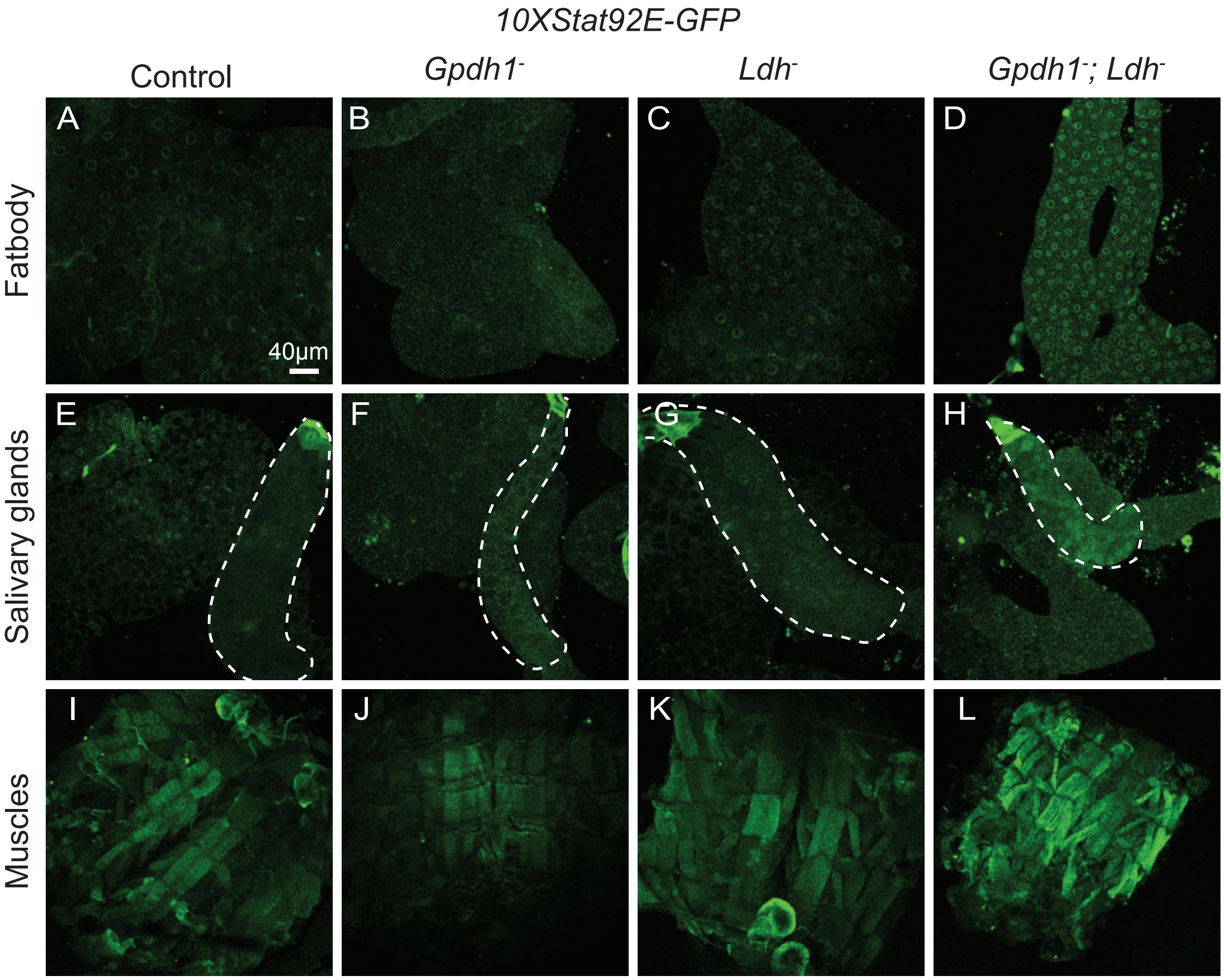


**Supplementary Figure 7. *Stat-GFP* expression is increased in larval tissues of *Gpdh1; Ldh* double mutants.** (A-L) Representative confocal images of (A-D) fat body, (E-H) salivary glands and (I-L) muscles showing *Stat-GFP* expression in control, $Gpdh1^{A10/B18}$, $Ldh^{16/17}$ and $Gpdh1^{A10/B18}$; $Ldh^{16/17}$ double mutants. The scale bar represents 40 μM. The scale bar in (A) applies to (B-L).
